# The Tricuspid Valve Maladapts in a Pulmonary Hypertension Rat Model

**DOI:** 10.64898/2026.07.08.737279

**Authors:** Chien-Yu Lin, Boguslaw Gaweda, Nikos Manthatis, Shreya Sreedhar, Vijay K. Dubey, Austin Goodyke, Tomasz A. Timek, Manuel K. Rausch

## Abstract

Tricuspid valve regurgitation is a frequent valve lesion and, if severe, an independent predictor of mortality. In most patients, the valve itself has historically been considered intact. Yet, we have previously shown that the valve may not be an innocent bystander. In multiple sheep models, we have shown that the tricuspid valve thickens and stiffens. This remodeling may contribute to valve disease. Our goal is to extend our investigation of tricuspid valve remodeling to a rodent model, potentially opening scientific opportunity and enabling scaling our studies.

To this end, we used pulmonary artery banding (PAB) in male rats to induce pressure overload and right ventricular remodeling. After excising the tricuspid valve, we quantified anterior leaflet morphology, mapped anterior leaflet thickness using optical coherence tomography, and evaluated anterior leaflet belly mechanics using a custom bulge testing system.

Compared with SHAM controls, PAB increased anterior leaflet area. Moreover, anterior leaflets in PAB animals exhibited region-specific thickening, with the largest increases near the annulus. Finally, anterior leaflets in PAB animals were significantly less compliant. However, leaflet stiffening stemmed from aforementioned thickening, i.e., structural stiffening, not constitutive stiffening.

Our findings demonstrate that we can reliably quantify leaflet area, thickness, and stiffness in the minuscule tricuspid valves of rats. We also show that tricuspid valve remodeling is not ovine-specific, but also affects the tricuspid valves of rats. Together, our findings support our hypothesis that tricuspid valves are not innocent bystanders in regurgitation, and that rats may serve as a scalable model system for future investigations.

**NEW & NOTEWORTHY:** Using a rat pulmonary artery banding model of pulmonary hypertension, we show that chronic right ventricular pressure overload induces leaflet enlargement and region-specific thickness remodeling of the tricuspid valve. Although structural mechanical metrics change under pressure loading, normalization by thickness reveals that geometric remodeling rather than intrinsic material stiffening predominates. These findings highlight leaflet structural (mal)adaptation as a potential contributor to functional tricuspid regurgitation and underscore the importance of considering leaflet geometry in therapeutic strategies.

## 1 INTRODUCTION

Tricuspid valve disease is extremely common. Approximately 1.6 million Americans suffer from moderate to severe leakage of the right atrioventricular heart valve [35, 8]. In most patients, tricuspid regurgitation (TR) is functional. In other words, it is secondary to other primary diseases, including left heart disease and pulmonary hypertension [7, 15]. Despite its secondary nature, TR has been shown to be an independent predictor of morbidity and mortality [27, 37, 39]. This somewhat recent recognition has led to a re-thinking of how we treat TR. In the past, the valve itself was seldom repaired or replaced. Instead, it was hoped that treatment of the primary disease, for example left-sided heart disease, would eventually lead to improvements in TR [3, 28]. However, today, treatment of the valve itself, especially in cases of severe TR, is more common [11, 40, 38].

As valve-specific treatment techniques, both surgical and interventional, have advanced, we have also made fundamental discoveries about the valve. Recent work has shown that tricuspid valve leaflets are not innocent bystanders but actively remodel in response to disease [34]. In particular, our group has demonstrated that the tricuspid valve remodels in two separate sheep models. Specifically, we showed that the tricuspid valve grows in area, thickens, and alters its stiffness [25, 17, 19]. In other words, the tricuspid valve maladapts. In our early discoveries, we hypothesized that these tissue changes could contribute to dysfunction and thus may provide a novel treatment target. In fact, in a high-fidelity computational model of the human tricuspid valve, we demonstrated that leaflet thickening and stiffening hinder coaptation [23].

However, much remains to be learned and understood about tricuspid valve remodeling. For example, the underlying stimuli are unclear. Moreover, we have yet to demonstrate that the detailed remodeling phenotypes we have observed are not limited to sheep. On the left side, similar remodeling phenomena have been demonstrated in the mitral valve across species, including humans [4, 5, 2, 6]; comparable cross-species evidence remains more limited for the tricuspid valve. Thus, the objective of this current study is to address this gap by testing for tricuspid valve remodeling in a rodent model. To this end, we use a surgical pulmonary hypertension model [9, 16, 26]. After eight weeks of disease progression, we explant the tricuspid valves and quantify leaflet area, leaflet thickness, and leaflet stiffness as markers of valvular remodeling.

## 2 MATERIALS AND METHODS

### 2.1 Pulmonary Hypertension Rat Model

All animals received humane care in compliance with the Principles of Laboratory Animal Care formulated by the National Society for Medical Research. The study protocol PROTO202400032 was approved by Michigan State University Institutional Animal Care and Use Committee. We housed and cared for animals at Michigan State University’s Grand Rapids Research Center animal facility.

Seven male Sprague-Dawley rats received a pulmonary artery band (PAB), which we applied using a half closed ligating clip around the main pulmonary artery. Additionally, seven sham animals (SHAM) underwent identical surgical and imaging procedures, excluding PAB. In brief, we pre-anesthetized 12 week-old animals with isoflurane in an induction chamber for 5-10 minutes (2.5% mix in 1.5L O_2_), intubated them with a 16Fr angiocatheter (BD), and placed them on a ventilator. During the procedure, we maintained anesthesia with a continuous infusion of 2% isoflurane with oxygen. Subsequently, we placed rats in a supine position on a heating pad and shaved the chest and right neck with a hair clipper. Performing a left thoracotomy, we opened the pericardium longitudinally. After identifying the left atrium, ascending aorta, and main pulmonary artery, we dissected the main pulmonary artery free from the ascending aorta. Subsequently, we exposed the right jugular vein from the clavicle halfway to the jaw angle, opened the jugular vein in its midportion, and advanced a pressure probe (Transonic Scisense Inc, London, ON, Canada) into the right ventricle (RV). We measured baseline RV systolic pressure and subsequently applied a half-closing ligating clip (Horizon® Titanium Medium Ligating Clip) with a pre-measured caliper applicator around the pulmonary artery to at least double the resting pressure in the RV. After achieving hemostasis, we closed the chest with three layers of running sutures.

Eight weeks after surgery, we euthanized the animals by CO_2_ asphyxiation and confirmed death with cervical dislocation. We then performed a sternotomy, excised the heart, opened the right and left atria, and removed the apex before flushing the tissue with 1 × PBS. Finally, we placed the heart in cryomedia consisting of DMEM:DMSO (9:1) supplemented with protease inhibitor (ThermoFisher, A32953, Waltham, MA) and slowly froze the tissue at -80 ^*o*^C for future testing.

### 2.2 Tissue Excision & Floating Morphology

Immediately before testing, we rapidly thawed the hearts. We accessed the right ventricle via the pulmonary artery, and opened the tricuspid annulus by incising the anterior-septal commissure of the tricuspid valve. We then isolated the tricuspid valve by dissecting the subvalvular apparatus (papillary muscles and chordae tendineae) from the ventricular wall and separating the leaflets along the annulus. Throughout the dissection, we rinsed the tissue with cold 1 × PBS supplemented with protease inhibitor (ThermoFisher, A32953, Waltham, MA) to preserve mechanical integrity. Next, we removed annular tissue as completely as possible. We then photographed the tricuspid valve leaflets floating in a thin layer of 1 × PBS over a calibrated grid for subsequent morphological analysis (Figure 1**a**). Using these images, we quantified the projected areas of the anterior leaflets and compared them between the SHAM and PAB groups.

**Figure 1.**
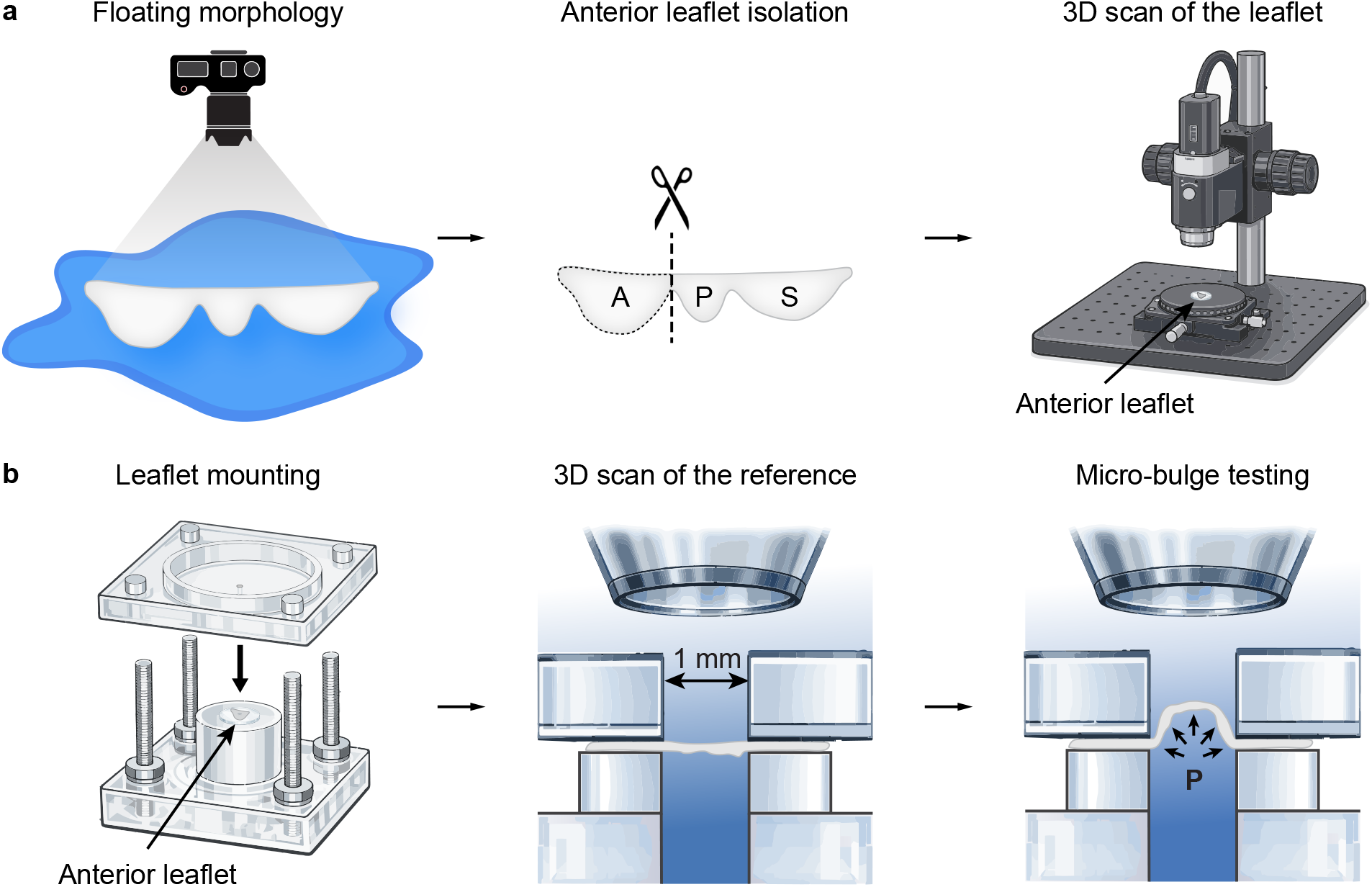
Experimental workflow for tricuspid valve leaflet morphology, optical coherence tomography (OCT)-based profilometry, and micro-bulge testing. **a**) After valve excision, we photographed the leaflets while floating on 1 × PBS for morphology analysis, then isolated the anterior leaflet and acquired a 3D OCT scan for tissue profilometry. **b**) We mounted the anterior leaflet over a circular aperture in the micro-bulge device, acquired a 3D OCT scan in a minimally pressurized reference configuration (*<*1 mmHg), and performed micro-bulge inflation while acquiring OCT B-scans to quantify pressure–deflection behavior. A, P, S denote anterior, posterior, and septal leaflets, respectively

### 2.3 Anterior Leaflet Profilometry

After photographing the tricuspid valve, we isolated the anterior leaflet for the tissue profilometry and the micro-bulge inflation tests described in the next section 2.4. Specifically, we trimmed away all chordae tendineae, and performed tissue profilometry using our optical coherence tomography (OCT) system (Telesto, Thorlabs, Inc., Newton, NJ, USA) with the 3D scan function in ThorImage OCT (v5.6.0). We placed a PDMS layer (Gel-Pak) on the system’s XY stage with a solid top plate (Item # OCT-XYR1, Thorlabs, Inc., Newton, NJ, USA), positioned the leaflet on the PDMS with the ventricular surface facing up in a relaxed configuration, and gently dabbed away excess 1 × PBS using absorbent tissue (KimWipes; Kimberly-Clark, Irving, TX, USA). We then acquired a 3D OCT scan of the leaflet (Figure 1**a**).

After imaging, we exported the 3D scans to ImageJ (v1.54g; National Institutes of Health, Bethesda, MD, USA) to convert the files to NRRD format. Next, we manually segmented each sample’s 3D volume in 3D Slicer (v5.8.1) [10] and exported the resulting segment to MATLAB (R2024a; MathWorks, Natick, MA, USA) in NIfTI (.nii) format. We then used a custom MATLAB program to partition the anterior leaflet thickness map into a 3 × 3 grid: three equidistant regions along the commissure-to-commissure direction between the anterior–posterior (AP) and anterior–septal (AS) commissures, and three equidistant regions from the near-annulus (NA) to the free edge (FE). We compared regional thickness values between the SHAM and PAB groups.

### 2.4 Micro-bulge Inflation Testing

After tissue profilometry, we performed micro-bulge inflation testing using our custom micro-bulge device described previously (see [24] for device construction and detailed protocols). We filled the chamber with room-temperature 1 × PBS and removed air bubbles before mounting each leaflet over a circular aperture in a PDMS gasket that defined the test region. We aligned the leaflet circumferential and radial directions with the device principal axes, mounted the top chamber, and controlled inflation using a syringe pump while measuring pressure with an in-line pressure transducer (Figure 1**b**). We preconditioned each sample with 10 loading cycles to 30 mmHg at a syringe displacement rate of 0.25 mL/min, then unmounted and remounted the tissue to remove pre-conditioning-induced slack. After remounting, we applied a minimal pressure preload (*<*1 mmHg) to enforce a consistent reference geometry without inducing measurable bulging, zeroed the pressure transducer, and acquired a 3D OCT scan in this reference state. We then inflated the tissue once to 30 mmHg while acquiring OCT B-scans in the circumferential and radial directions throughout loading (5 frames/s).

We exported and segmented the 3D reference-geometry scan of the test region as described for anterior leaflet profilometry (Section 2.3) to obtain the mean thickness of the test region. Then, we exported the OCT B-scans acquired during micro-bulge inflation to MATLAB, tracked the maximum (approximately central) out-of-plane deflection *w* in each imaging plane over the loading history, and paired deflection with the recorded pressure *p* to generate pressure–deflection curves for each sample. We first quantified structural stiffness directly from the pressure–deflection response for comparisons between the SHAM and PAB groups. Specifically, we defined toe stiffness and calf stiffness as the slopes of tangent lines to the pressure–deflection curve in the lowand high-deflection regimes, respectively. To obtain these tangents, we fit a toe tangent line and a calf tangent line by linear regression in a small neighborhood (ten data pairs) around *p* = 0 mmHg and *p* = 20 mmHg. We defined the transition deflection *w*_*T*_ as the deflection value corresponding to the data point on the pressure–deflection curve that was closest to the intersection of the toe and calf tangent lines. We then used the mean thickness together with the pressure–deflection data to compute thickness-normalized stress–strain curves, from which we calculated tangent moduli and the transition strain for group comparisons, as discussed in the following paragraphs.

Following commonly adopted bulge-test analyses [32, 33, 36, 42], we assumed the deformed membrane formed a spherical cap, which allowed us to describe the stress state using membrane equilibrium relations. Under this assumption, the equibiaxial membrane stress (*σ*) is

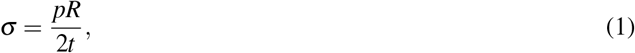

where *R* is the radius of curvature of the spherical cap and *t* is the membrane thickness.

Let *a* be the radius of the circular aperture. From spherical-cap geometry, we expressed the radius of curvature in terms of the measured central deflection *w* as

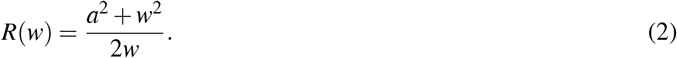

Substituting this expression into the stress relation yields

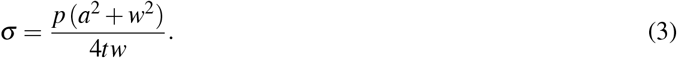

To characterize deformation, we defined the stretch as

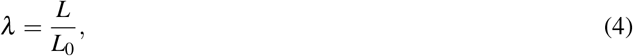

where *L*_0_ is the reference length, and *L* is the deformed arc length along the spherical surface from the membrane center to the clamped boundary.

For the assumed spherical geometry, the arc length is

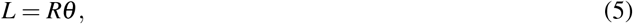

where the central angle satisfies

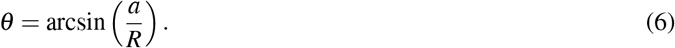

Since the reference length corresponds to the aperture radius (*L*_0_ = *a*), the stretch becomes

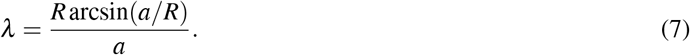

Since the bulge-test deformations were not infinitesimal, we did not use small-strain approximations and instead quantified deformation using a finite-strain measure. We computed the Green–Lagrange strain from the stretch ratio, which remains valid for large deformations while referencing the initial configuration, as

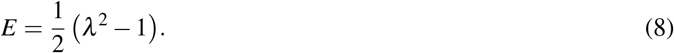

We shifted all stress–strain curves such that both stress and strain were zero at zero applied pressure to enforce a consistent reference configuration across samples.

To quantify the evolution of material stiffness with increasing pressurization, we defined the tangent modulus as the local slope of the stress–strain curve,

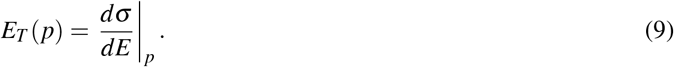

Similar to our structural-stiffness analysis, we evaluated *E*_*T*_ at two prescribed pressure levels: *p* = 0 mmHg (toe modulus) and *p* = 20 mmHg (calf modulus), using the same fitting procedure described above. Finally, we defined the transition strain as the strain value corresponding to the data point on the stress–strain curve that was closest to the intersection of the toe and calf-region tangent lines.

### 2.5 Statistics

We conducted all statistical analyses in R (Version 4.5.2), and we defined statistical significance as *p <* 0.05. For the analysis of regional thickness maps, we fit linear mixed-effects models using the R package *afex* (Version 1.4.1) to account for repeated measurements within each leaflet and performed post hoc comparisons using Tukey-adjusted tests. For outcomes that required only a single between-group comparison, we used Welch’s t-tests. Unless indicated otherwise, we report data as mean ± 1 Standard Deviation.

## 3 RESULTS

### 3.1 Pulmonary artery banding increased systolic pulmonary artery pressure in rats

Baseline systolic pulmonary artery pressure (PAP) was similar between sham (SHAM) and pulmonary artery banding (PAB) animals (Table 1). In PAB animals, systolic PAP increased from 19.7 ± 1.0 mmHg before clip placement to 39.6 ± 2.6 mmHg after clip placement (*p <* 0.001), representing an approximately 2-fold increase and confirming successful pulmonary artery constriction.

**Table 1.**
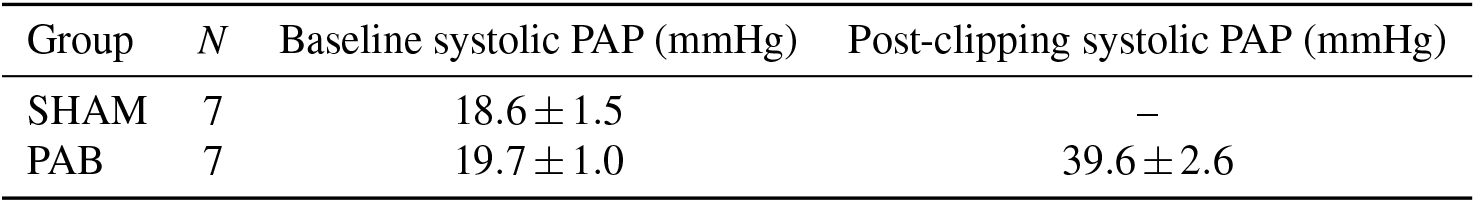
Systolic pulmonary artery pressure (PAP) measurements in sham (SHAM) and pulmonary artery banding (PAB) animals. SHAM animals were measured at baseline only, whereas PAB animals were measured before and after clip placement, demonstrating the acute increase in PAP following clipping

| Group | *N* | Baseline systolic PAP (mmHg) | Post-clipping systolic PAP (mmHg) |
| --- | --- | --- | --- |
| SHAM | 7 | $18.6 \pm 1.5$ | — |
| PAB | 7 | $19.7 \pm 1.0$ | $39.6 \pm 2.6$ |

### 3.2 Pulmonary artery banding increased anterior tricuspid leaflet area

Figure 2 shows that pulmonary artery banding (PAB) increased anterior tricuspid leaflet area compared to SHAM (*N* = 7). Anterior leaflet area increased from 11.8 ± 3.2 mm^2^ in SHAM to 14.9 ± 1.2 mm^2^ in PAB, an increase of approximately 27% (Welch’s *t*-test, p = 0.042; Figure 2).

**Figure 2.**
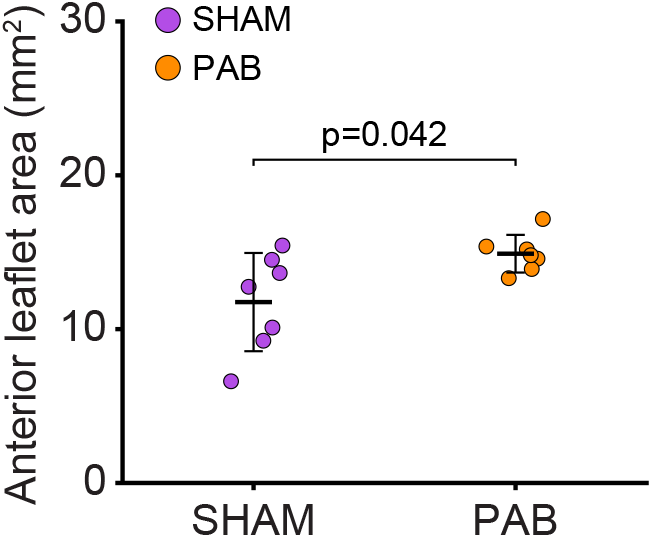
Anterior tricuspid leaflet area in sham (SHAM) and pulmonary artery banding (PAB) animals (*N* = 7). PAB animals exhibited significantly larger anterior leaflet area than SHAM animals

### 3.3 Pulmonary artery banding increased tricuspid leaflet thickness with regional heterogeneity

Thickness profilometry revealed both baseline spatial heterogeneity and disease-associated thickening in PAB leaflets (Figure 3). Representative anterior leaflet thickness maps illustrated visually higher thickness in PAB compared to SHAM, with the largest values concentrated near the annulus (Figure 3**a**). To quantify regional differences, we segmented each leaflet into a 3 × 3 grid spanning the radial direction (near-annulus, belly, free edge) and the circumferential direction (anterior–posterior commissure, middle, anterior–septal commissure), then computed mean thickness for each region (Figure 3**b**; *N* = 6). Across regions, the absolute thickness difference map (PAB – SHAM) showed positive differences throughout the leaflet, indicating thicker tissue in PAB (Figure 3**c**, top). The fold-change map (PAB/SHAM) further highlighted regional variation in the magnitude of thickening (Figure 3**c**, bottom; *N* = 6).

**Figure 3.**
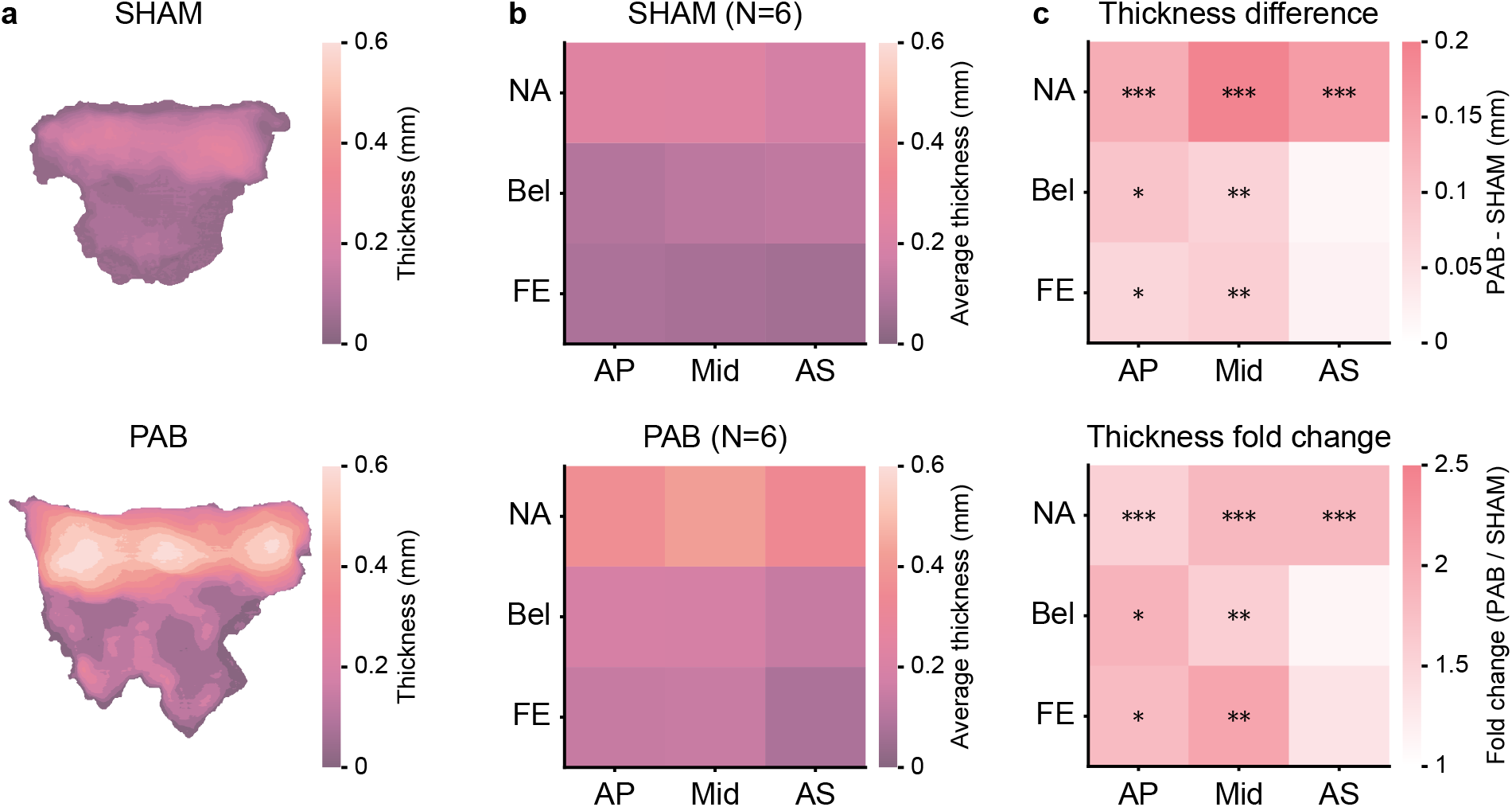
PAB leaflets were thicker than SHAM leaflets in a region-dependent manner. **a**) Representative thickness maps of anterior leaflets from PAB and SHAM groups. **b**) All samples in the SHAM and PAB groups were segmented into 3 × 3 meshes for thickness comparison. The average thickness within each region is shown for SHAM and PAB groups (*N* = 6). **c**) Absolute thickness difference (PAB - SHAM) and fold change (PAB / SHAM) maps illustrated regional variation between groups. Significant differences between SHAM and PAB are indicated by * (*p <* 0.05), ** (*p <* 0.01), and *** (*p <* 0.001). NA: Near-annulus, Bel: Belly, FE: Free edge, AP: Anterior–posterior commissure, Mid: Middle, AS: Anterior–septal commissure

### 3.4 Pressure–deflection analysis of the micro-bulge tests showed increased toe stiffness in PAB leaflets

Pressure–deflection behavior of anterior tricuspid valve leaflets differed between SHAM and PAB groups (Figure 4; *N* = 6). Across the measured pressure range, PAB specimens exhibited reduced maximum deflection relative to SHAM, consistent with reduced structural compliance (Figure 4**a**). To quantify these differences, we extracted three metrics from the pressure–deflection response: toe stiffness, calf stiffness, and the transition deflection (*w*_*T*_) (Figure 4**b**). Toe stiffness was significantly higher in PAB than SHAM (*p* = 0.012; Figure 4**c**, left). In contrast, calf stiffness did not differ between groups (Figure 4**c**, middle). Transition deflection showed the same trend (lower *w*_*T*_ in PAB), but this difference was not statistically significant (*p* = 0.086; Figure 4**c**, right).

**Figure 4.**
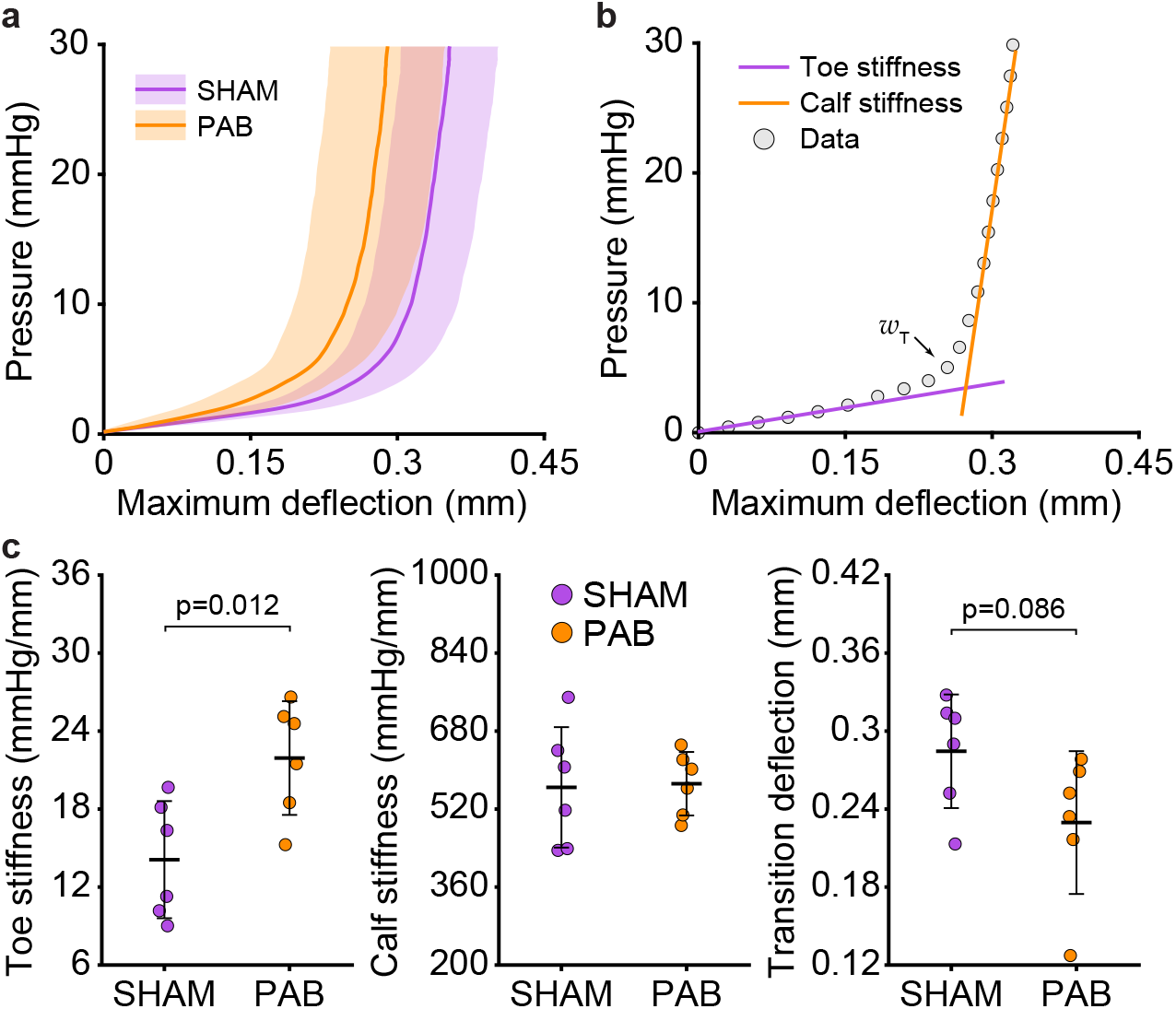
Pressure–deflection analysis of the micro-bulge tests revealed increased toe stiffness in PAB leaflets, indicating reduced compliance in remodeled tissues; transition deflection demonstrated the same pattern, though without statistical significance (*N* = 6). **a**) Pressure-deflection curves for SHAM and PAB groups. **b**) Explanation of three quantitative metrics of the tissue structural mechanics: Toe stiffness, Calf stiffness, and Transition deflection (*w*_T_). **c**) Quantitative comparison of structural mechanics in anterior tricuspid valve leaflets from SHAM and PAB groups

### 3.5 Thickness normalization attenuated toe-region stiffening in stress–strain analysis

We next converted the micro-bulge pressure–deflection data to a stress–strain response using a spherical-cap assumption and computed a thickness-normalized tangent modulus as a function of pressure (Figure 5**a**; *N* = 6). The average thickness measured within the bulge-test region was higher in PAB than SHAM (Figure 5**b**). After thickness normalization, toe modulus did not differ between PAB and SHAM (Figure 5**c**, left). Similarly, calf modulus did not differ between groups (Figure 5**c**, middle). Transition strain exhibited a trend toward greater compliance in SHAM, though this difference did not reach statistical significance (*p* = 0.057; Figure 5**c**, right).

**Figure 5.**
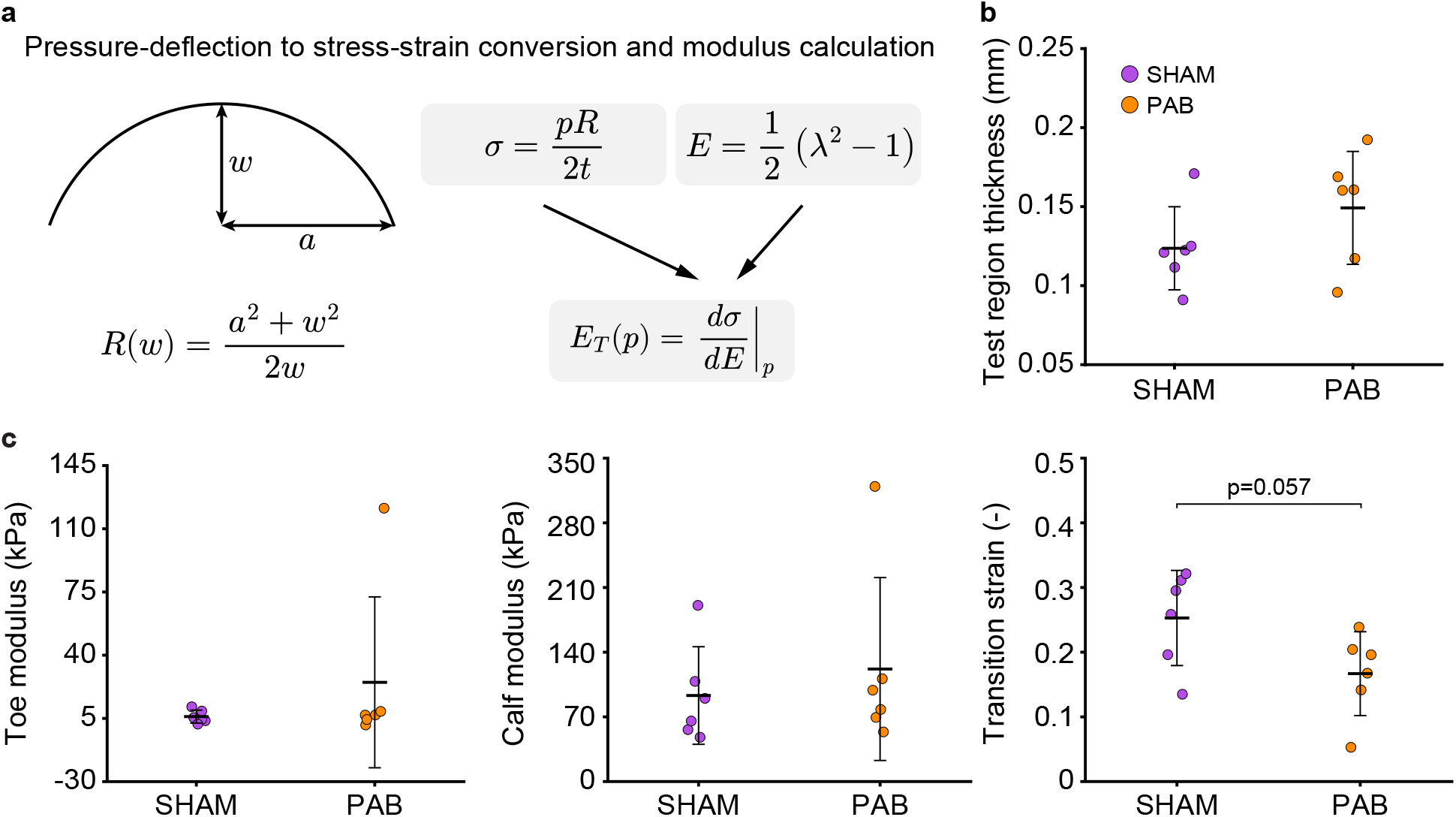
Stress–strain analysis derived from micro-bulge testing demonstrated that thickness normalization attenuated the toe-region stiffening observed in pressure-deflection analysis, suggesting that thickening contributes to mechanical differences between groups (*N* = 6). Transition strain exhibited a trend toward greater compliance in SHAM leaflets, though without statistical significance. **a**) Conversion of pressure-deflection data to stress-strain response and thickness-normalized tangent modulus. Assuming the bulged membrane forms a spherical cap with aperture radius *a* and central deflection *w*, the corresponding radius of curvature *R*(*w*) is calculated from this geometry and used to determine membrane stress, strain, and the tangent modulus. Please see Section 2.4 for details. **b**) Comparison of average test-region thickness between SHAM and PAB groups. **c**) Quantitative comparison of thickness-normalized toe modulus, calf modulus, and transition strain (*E*_T_) in anterior tricuspid valve leaflets from SHAM and PAB groups

## 4 DISCUSSION

In this study, we set out to test whether the tricuspid valve in a small-animal tricuspid regurgitation (TR) model undergoes leaflet-level remodeling analogous to that observed in our ovine studies. In rats subjected to pulmonary artery banding (PAB), we found clear evidence of anterior tricuspid leaflet maladaptation: anterior leaflet area increased, thickness increased with marked spatial heterogeneity, and structural stiffness increased in the low-deflection (toe) regime. Together, these findings show that chronic right-sided pressure overload is sufficient to drive geometric and structural mechanical remodeling of the tricuspid valve in a rodent model, supporting the concept that tricuspid leaflet maladaptation is not limited to large animals and providing a cross-species extension of our prior ovine work.

One key component of this maladaptive response is leaflet enlargement. In this study, PAB increased anterior tricuspid leaflet area versus SHAM, indicating leaflet-level geometric remodeling of the tricuspid valve in this rodent pulmonary hypertension model. This finding aligns with atrioventricular valve literature showing that chronic alterations in loading can be accompanied by progressive leaflet enlargement, and that enlargement can reflect an active remodeling response rather than purely passive distension. In an ovine mitral tethering model designed to isolate mechanical stretch, diastolic leaflet area increased over time and coincided with substantial leaflet thickening and cellular/matrix changes, consistent with stretch-triggered remodeling [5]. Evidence for stretch-driven leaflet growth also comes from work quantifying leaflet area increases attributable to both circumferential and radial enlargement, supporting the concept that sustained mechanical stretch can provide a stimulus for leaflet growth [29].

The observed increase in anterior leaflet area suggests that tricuspid leaflet remodeling is an important component of the valve response to chronic right-sided pressure overload. Such remodeling is unlikely to be uniform because each leaflet experiences distinct loading and boundary conditions, which can drive different adaptive responses. This concept is also supported by our prior large-animal tachycardia-induced cardiomyopathy model, in which anterior and septal leaflet areas increased significantly, whereas posterior leaflet area remained unchanged [19]. Importantly, prior tricuspid pressure-overload work indicates that leaflet growth can co-occur with other remodeling modes (for example, thickening and stiffness changes) rather than proceeding as isolated geometric adaptation [17]. Clinical pulmonary hypertension data further support the functional relevance of leaflet remodeling: although tricuspid leaflet area increases in pulmonary hypertension, this adaptation is often insufficient relative to the systolic closure area, resulting in an inadequate leaflet-area-to-closure-area ratio in severe tricuspid regurgitation [1]. Although we did not quantify leaflet strains or mechanobiological pathways directly, the observed anterior leaflet enlargement defines a clear remodeling phenotype in this rodent model and motivates further investigation into how the local mechanical environment shapes tricuspid leaflet remodeling.

Beyond leaflet area, PAB also induced significant leaflet thickening with pronounced spatial heterogeneity. Thickness maps and 3 × 3 regional averaging show that thickening is not uniform across the leaflet surface; instead, the intrinsic radial gradient (thicker near the annulus and thinner toward the free edge) is preserved with superimposed PAB-associated increases. These thickness changes are mechanically meaningful because structural behavior under pressurization depends on geometry as well as tissue material properties; increased thickness can reduce deflection and shift the apparent stiffness of the pressure-deflection response even if intrinsic moduli are unchanged, and ignoring spatial thickness heterogeneity can substantially bias biomechanical predictions [21]. Consistent with this interpretation, large-animal pressure-overload studies report leaflet thickening as part of functional tricuspid regurgitation remodeling, often alongside other tissue-level changes [17]. Together, anterior leaflet enlargement and spatially patterned thickening suggest that tricuspid valve remodeling in this rodent model involves both whole-leaflet geometric adaptation and regionally heterogeneous structural remodeling.

The micro-bulge tests further demonstrated altered *structural* mechanics in PAB, characterized by reduced deflection under pressure and a significant increase in toe-region stiffness. Notably, calf stiffness did not differ between groups, and transition deflection showed a similar (lower) trend in PAB but did not reach statistical significance. This pattern is consistent with remodeling that predominantly affects the low-deflection regime of the structural response, where geometry (including thickness) and early recruitment of load-bearing architecture exert strong influence, while comparatively preserving the high-deflection regime [31, 30].

When the same micro-bulge data were converted to a thickness-normalized stress–strain response, between-group differences were markedly reduced. Specifically, PAB specimens were thicker within the bulge-test region, and after accounting for this increased thickness, toe modulus did not differ significantly between groups. Considered alongside the pressure–deflection results, these findings support the interpretation that the elevated toe-region *structural* stiffness in PAB is driven, at least in part, by thickening within the tested region: thickness normalization removes a major geometric contribution, thereby diminishing the apparent between-group difference. More broadly, the results highlight a key interpretive point for valve remodeling studies: structural measurements integrate geometry and material behavior, and geometric remodeling (especially thickening) can manifest as apparent stiffening even when intrinsic material moduli are unchanged or only modestly altered.

This geometry-versus-material distinction also helps relate our prior sheep findings across disease models. In large animals, leaflet area and thickness often show clearer and more consistent remodeling signals than stiffness, whereas stiffness changes may emerge only in certain leaflets, directions, or curve regions (toe versus calf) rather than as a uniform increase across the valve [19, 17]. Our rodent PAB results align with this pattern, showing prominent geometric remodeling (increased anterior leaflet area and thickness) while intrinsic material changes remain subtle, specifically concentrated in the toe regime and attenuated after thickness normalization. Quantifying such native leaflet maladaptation in pressure overload may complement device studies showing how shape/size profoundly affect post-repair mechanics [14]. This native remodeling information can help clinicians refine device selection to match disease-specific phenotypes.

To translate these findings into targeted treatments, an important next step is to determine what biological pathways drive leaflet growth and thickening in pressure-overload tricuspid disease. Recent large-animal pulmonary artery banding studies in sheep have shown that tricuspid leaflet remodeling is accompanied by marked changes in gene expression and activation of specific molecular pathways within the leaflet tissue [12, 13], underscoring that leaflet maladaptation is a biologically regulated process. In light of this, the genetic tractability of rodent models provides an opportunity to link specific molecular and cellular mechanisms to leaflet maladaptation and to explore strategies aimed at modifying leaflet remodeling in pulmonary hypertension.

In addition to enabling mechanistic studies, the rodent PAB model provides an opportunity to scale future investigations of tricuspid leaflet remodeling. Compared with large-animal models, rodent studies can support larger cohort sizes, more efficient evaluation across disease time points, and faster testing of candidate pathways or interventions [18]. This scalability is especially important for defining the temporal sequence of leaflet growth, thickening, and mechanical change during pressure-overload disease progression. Moreover, pulmonary artery banding has been widely used as an experimental model of right-heart pressure overload and provides a useful framework for mechanistic and interventional studies [22]. With access to genetically modified rodents, future studies can move beyond phenotypic characterization to directly test the roles of specific genes, cell populations, and signaling pathways in driving tricuspid valve remodeling [41]. In this way, the present model provides not only a cross-species extension of leaflet maladaptation, but also an opportunity for mechanistic studies of tricuspid valve disease.

A key limitation of the present mechanical testing is that our mounting strategy restricts micro-bulge measurements to the leaflet belly region and, in practice, to the anterior leaflet. This is particularly relevant given the pronounced regional heterogeneity observed in our thickness maps, which suggests that the largest remodeling and thus the largest material property differences may occur outside the belly region interrogated by bulge testing (for example, closer to the annulus or toward the free edge). Consistent with this concern, regional biaxial testing in porcine atrioventricular valve leaflets has demonstrated substantial spatial variation in leaflet mechanics across regions [20]. Accordingly, future approaches that enable spatially resolved mechanical quantification, such as nanoindentation, may reveal localized mechanical phenotypes that are not captured by a single belly-region measurement.

In conclusion, we quantified a cohesive tricuspid leaflet remodeling signature in the PAB model, combining geometric changes (increased anterior leaflet area and heterogeneous thickening) with mechanical changes (structural toe-stiffening with attenuated thickness-normalized modulus differences). These findings demonstrate that chronic right-heart pressure overload in rats is sufficient to induce leaflet-level tricuspid valve maladaptation and provide a foundation for targeted studies of the biological mechanisms that drive leaflet growth and thickening in pressure-overload tricuspid disease.

## 5 DATA AVAILABILITY

The data supporting the findings of this study will be deposited in our Data Repository and will be made publicly available upon publication.

## 6 GRANTS

This work was partially supported by an NIH grant to MKR and TAT (R01HL165251), an AHA Predoctoral Fellowship to CYL (25PRE1363276), an NIH T32 Training Fellowship to SS (T32 EB007507), and a grant from the DeVos Cardiovascular Research Program.

## 7 DISCLOSURES

The authors declare the following financial interests/personal relationships which may be considered as potential competing interests: Manuel K. Rausch has a speaking agreement with Edwards Lifesciences. None of the other authors have conflicts to declare.

## 8 DISCLAIMERS

The content is solely the responsibility of the authors and does not necessarily represent the official views of the National Heart, Lung, and Blood Institute of the National Institutes of Health.

## 9 AUTHOR CONTRIBUTIONS

CYL, BG, MKR, and TAT conceived and designed research; CYL, BG, and AG performed experiments, CYL, NM, and VKD analyzed data, CYL, NM, and MKR interpreted results of experiments, CYL prepared figures, CYL, NM, and MKR drafted manuscript, CYL, BG, SS, VKD, MKR edited and revised manuscript, CYL, BG, NM, SS, VKD, AG, TAT, and MKR approved final version of manuscript.

